# Substantial somatic genomic variation and selection for *BCOR* mutations in human induced pluripotent stem cells

**DOI:** 10.1101/2021.02.04.429731

**Authors:** Foad J Rouhani, Xueqing Zou, Petr Danecek, Tauanne Dias Amarante, Gene Koh, Qianxin Wu, Yasin Memari, Richard Durbin, Inigo Martincorena, Andrew R Bassett, Daniel Gaffney, Serena Nik-Zainal

## Abstract

Human Induced Pluripotent Stem Cells (hiPSC) are an established patient-specific model system where opportunities are emerging for cell-based therapies. We contrast hiPSCs derived from different tissues, skin and blood, in the same individual. We show extensive single-nucleotide mutagenesis in all hiPSC lines, although fibroblast-derived hiPSCs (F-hiPSCs) are particularly heavily mutagenized by ultraviolet(UV)-related damage. We utilize genome sequencing data on 454 F-hiPSCs and 44 blood-derived hiPSCs (B-hiPSCs) to gain further insights. Across 324 whole genome sequenced(WGS) F-hiPSCs derived by the Human Induced Pluripotent Stem Cell Initiative (HipSci), UV-related damage is present in ~72% of cell lines, sometimes causing substantial mutagenesis (range 0.25-15 per Mb). Furthermore, we find remarkable genomic heterogeneity between independent F-hiPSC clones derived from the same reprogramming process in the same donor, due to oligoclonal populations within fibroblasts. Combining WGS and exome-sequencing data of 452 HipSci F-hiPSCs, we identify 272 predicted pathogenic mutations in cancer-related genes, of which 21 genes were hit recurrently three or more times, involving 77 (17%) lines. Notably, 151 of 272 mutations were present in starting fibroblast populations suggesting that more than half of putative driver events in F-hiPSCs were acquired *in vivo*. In contrast, B-hiPSCs reprogrammed from erythroblasts show lower levels of genome-wide mutations (range 0.28-1.4 per Mb), no UV damage, but a strikingly high prevalence of acquired *BCOR* mutations of ~57%, indicative of strong selection pressure. All hiPSCs had otherwise stable, diploid genomes on karyotypic pre-screening, highlighting how copy-number-based approaches do not have the required resolution to detect widespread nucleotide mutagenesis. This work strongly suggests that models for cell-based therapies require detailed nucleotide-resolution characterization prior to clinical application.

## Main text

Yamanaka demonstrated that over-expression of particular transcription factors in somatic cells could produce hiPSCs^1^. In regenerative medicine, hiPSCs and latterly, organoids, have become attractive model systems because they can be propagated indefinitely and differentiated into many cell types. Specifically, hiPSCs have been adopted as a cellular model of choice for disease modelling and cell-based therapies^2,3^.

Some work has been done to explore genomic integrity and tumorigenic potential of human pluripotent stem cells^4–8^. However, systematic large-scale, whole genome assessments of mutagenesis at single-nucleotide resolution have been limited^4–8^. Cultured human embryonic stem cells (hESCs) have been reported to harbor recurrent chromosomal-scale genomic abnormalities ascribed to selection pressure^9–13^. Moreover, a study across 140 hESCs identified five subclonal dominant-negative *TP53* mutations. Using RNA-seq data from another 117 pluripotent cell lines, the authors reported nine additional *TP53* mutations^14^. The authors postulated that this could be due to selection conferred by prototypical *TP53* driver mutations. Yet, a subsequent study showed a low mutation burden in clinical grade hESCs and no mutations in *TP53* or other survival-promoting genes^15^.

The mutational burden in any given hiPSC comprises mutations that were pre-existing in the parental somatic cells from which it was derived, and mutations which have accumulated over the course of reprogramming, cell culture and passaging^7,16–22^. Several small-scale genomic studies have shown that in some cell lines, pre-existing somatic mutations make up a significant proportion of the total burden^22–28^. With the advent of clinical trials using hiPSCs (e.g. NCT03815071, NCT04339764, R000038278 and R000013279), comes the need to gain an in-depth understanding of the mutational landscape and potential risk of using these cells^29,30^. Here, we contrast hiPSCs from one individual, derived from two of the most commonly utilized tissues for hiPSC derivation (skin and blood). We then comprehensively assess a wide array of hiPSCs, including lines from one of the world’s largest stem cell banks, HipSci, and an alternative cohort called Insignia. All lines have been karyotypically pre-screened and believed to be chromosomally stable. We utilize combinations of whole genome (WGS) and whole exome (WES) sequencing of nearly 500 hiPSC samples to understand the extent of genomic damage in this model system and their possible implications.

### F-hiPSCs have greater levels of genome-wide substitutions and indels than B-hiPSCs

To understand whether the source of somatic cells used to make hiPSCs impacted on mutational load, we compared genomic variation in two independent skin-derived (F-hiPSCs) and two independent blood-derived hiPSCs (B-hiPSCs) from a 22-year-old healthy adult male (S2) (Fig. 1A). F-hiPSCs were derived from skin fibroblasts and B-hiPSCs were derived from peripheral blood Endothelial Progenitor Cells (EPCs). Additionally, we derived F-hiPSCs and B-hiPSCs from six healthy males (S7, oaqd, paab, yemz, qorq, quls) and four healthy females (iudw, laey, eipl, fawm) (Fig. 1A). All hiPSC clones were subjected to WGS and germline variants removed to reveal mutations acquired *in vivo* as a somatic cell and *in vitro* following reprogramming and cell culture.

**Figure 1.**
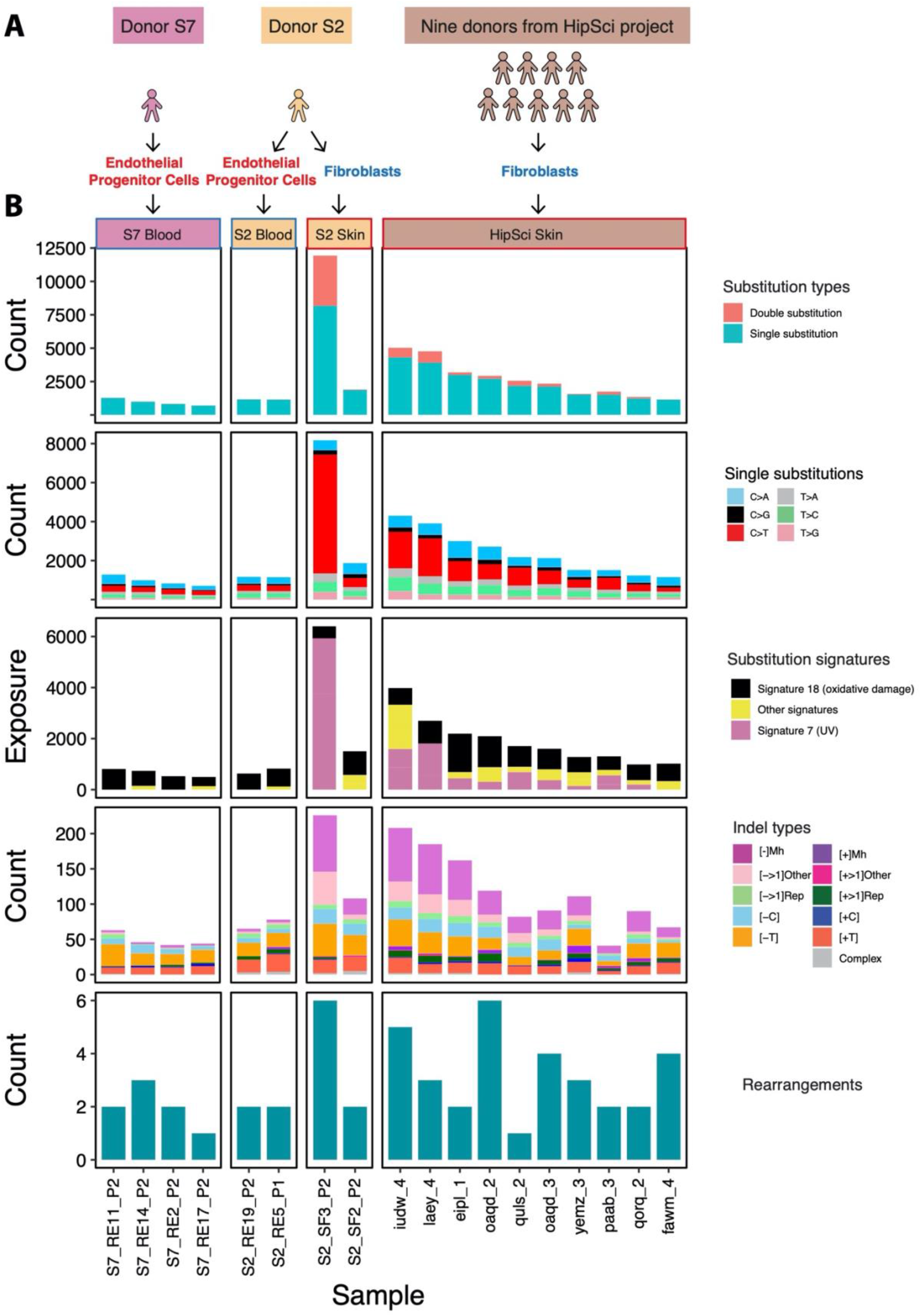
Mutation burden in blood-derived and fibroblast-derived hiPSCs. (A) Source of hiPSCs. Multiple hiPSC lines created from patient S2 contrasted to skin- and blood-derived hiPSCs created from ten other individuals. (B) Mutation burden of substitutions, double substitutions (first row), substitution types (second row), skin-derived signatures (third row), indel types (fourth row), as well as rearrangements (lowest panels). UV-specific features, e.g., elevated CC>TT double substitutions and UV mutational signatures were enriched in skin-derived hiPSCs.

WGS analysis revealed a greater number of mutations (~4.4 increase) in F-hiPSCs, as compared to B-hiPSCs, in individual S2 and in lines derived from the other ten donors (Fig. 1B and Table S2). There were very few structural variants (SVs) observed in these lines; thus, chromosomal-scale aberrations were not distinguishing between the F-hiPSCs and B-hiPSCs (Table S2). We noted considerable heterogeneity in total numbers of mutations between sister hiPSCs from the same donor, S2: one F-hiPSC line (S2_SF3_P2) had 8171 single substitutions, 1879 double substitutions and 226 indels, whereas the other F-hiPSC line (S2_SF2_P2) had 1,873 single substitutions, 17 double substitutions and 71 indels (Fig. 1B). Mutational signature analysis demonstrated striking predominance of UV-associated substitutions (Reference Signature/COSMIC) Signature 7^31^ in most of the F-hiPSCs, characterized by C>T transitions at T<u>C</u>A, T<u>C</u>C and T<u>C</u>T (Fig. 1B and Fig. S1). This is consistent with previously published work which has attributed UV signatures in hiPSCs to pre-existing damage in skin fibroblasts^8^. In contrast, B-hiPSCs did not show any evidence of UV damage but showed patterns consistent with possible oxidative damage (Signature 18, characterized by C>A mutations at T<u>C</u>T, G<u>C</u>A and A<u>C</u>A, Fig. 1 and Fig. S1). Consistent with *in vitro* studies^32,33^, double substitutions were enriched in UV-damaged F-hiPSCs (Fig. 1B). In all, we concluded that F-hiPSCs carry UV-related genomic damage as a result of sunlight exposure *in vivo* that does not manifest in B-hiPSCs. Importantly, copy-number-based screening for CNAs underestimated the substantial substitution/indel-based variation that exists in hiPSCs.

### High prevalence of UV-associated DNA damage in F-hiPSCs

We asked whether these findings were applicable across F-hiPSC lines generally. Therefore, we interrogated all lines in the HipSci stem cell bank, comprising 452 F-hiPSCs generated from 288 healthy individuals (Fig. 2A). These F-hiPSC lines were generated using Sendai virus, cultured on irradiated mouse embryonic fibroblast feeder cells which have been reported to help reduce genetic instability^20^ and were expanded prior to sequencing (range 7-46 passages, median18 passages, Fig. S2)^34^. WGS data were available for 324 F-hiPSC and matched fibroblast lines (Fig. 2A). We sought somatic mutations and identified 1,359,098 substitutions, 135,299 CC>TT double substitutions and 54,075 indels. Large variations in mutation distribution were noted: 692 – 37,120 substitutions, 0 – 7,864 CC>TT double substitutions and 17 – 641 indels per sample detected respectively (Fig. 2B).

**Figure 2.**
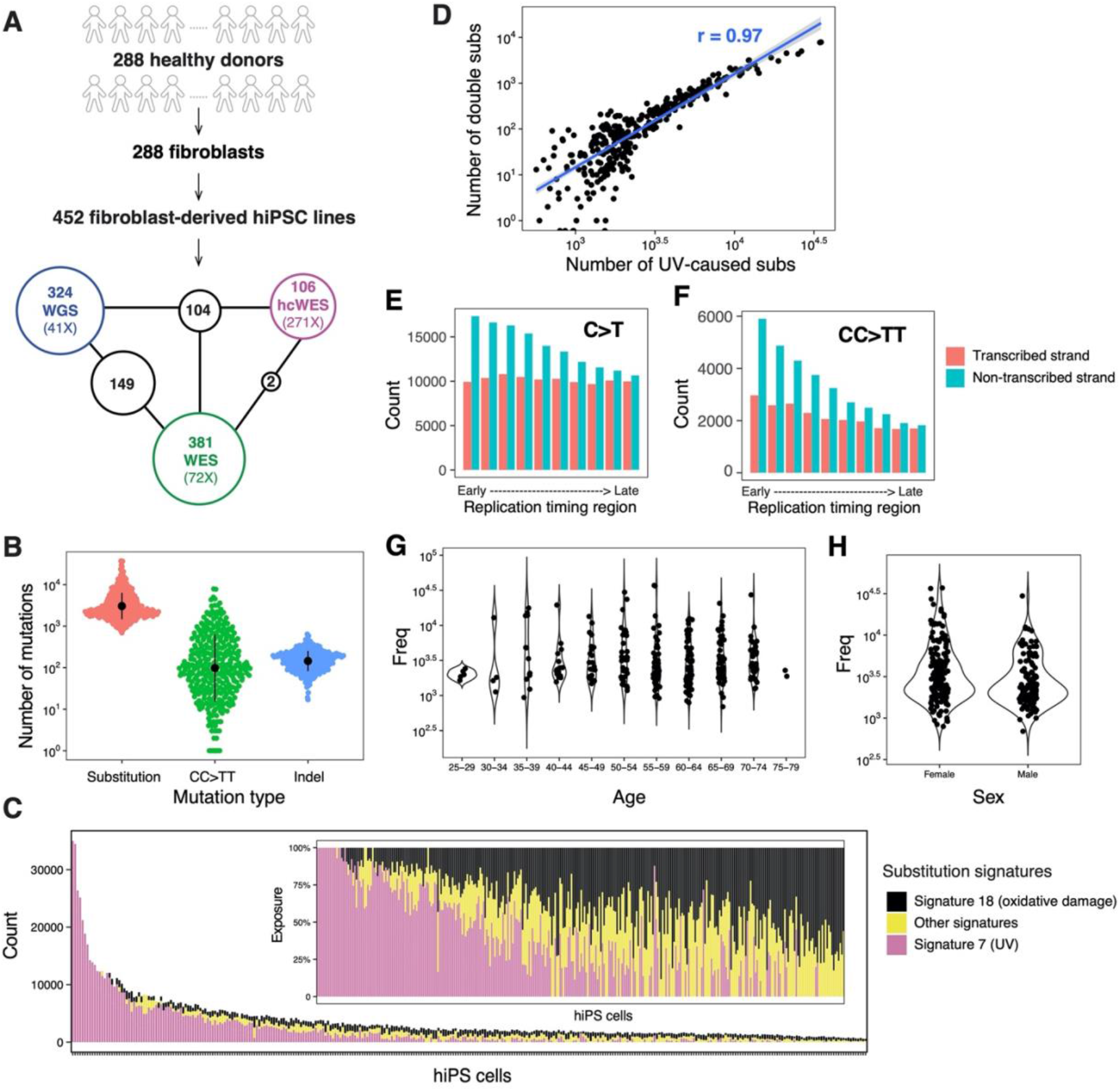
Mutation burden and mutational signatures in fibroblast-derived hiPSCs. (A) Summary of data. 452 fibroblast-derived hiPSC lines were generated from 288 healthy donors. 324 hiPSCs were whole-genome sequenced (WGS) (coverage: 41X), 381 were whole-exome sequenced (WES) (coverage: 72X) and 106 were WES with high coverage (271X). Table S3 provides source information. Numbers within black circles denote the number of samples that had data from multiple sequencing experiments. (B) Mutation burden of substitutions, CC>TT double substitutions and indels in fibroblast-derived iPSCs from WGS. Data summary is provided in Table S4. Black dots and error represent mean ± SD of hiPSC observations. (C) Distribution of mutational signatures in 324 fibroblast-derived iPSC lines. The inset figure shows the relative exposures of mutational signatures types. (D) Relationships between mutation burdens of CC>TT double substitutions and UV-caused mutation burden of substitutions in fibroblast-derived hiPSCs. (E)(F) Histogram of aggregated mutation burden on transcribed (red) and non-transcribed (cyan) strands for C>T (E) and CC>TT (F). Transcriptional strand asymmetry across replication timing regions was observed. (G)(H) Distribution of substitution burden of hiPSCs with respect to donor’s age (G) and gender (H).

Mutational signature analysis^31^ revealed that 72% of F-hiPSCs carried detecTable Substitution signatures of UV damage (Fig. 2C). hiPSCs with greater quantities of UV-associated substitution signatures showed strong positive correlations with UV-associated CC>TT double substitutions (Fig. 2D), microhomology-mediated deletions (mh-deletions) (Fig. S3) and C/T indels at short repeats (p value < 0.0001) (Fig. S3). A clear transcriptional strand bias with an excess of C>T and CC>TT on the non-transcribed strand was observed and was more striking in early replication timing domains than in late ones (Fig. 2E and 2F). These findings are consistent with previous reports of UV-related mutagenesis including the more efficient activity of transcription-coupled nucleotide excision repair (TC-NER) particularly in gene-rich early replication timing regions^35–38^. Of note, there was no correlation between mutation burden of F-hiPSC and donor age or gender (Fig. 2G and 2H).

### Substantial genomic heterogeneity between F-hiPSCs is due to clonal populations present in the starting material

F-hiPSCs comprise the majority of hiPSCs in stem cell banks globally and are a prime candidate for use in disease modelling and cell-based therapies. Yet, we and others5,6,23 observed substantial heterogeneity between hiPSC sister lines (Fig. 1, subject S2). It has been postulated that this heterogeneity may result from the presence of clones within the fibroblast population. To explore this in more detail, we compared mutational profiles of 118 pairs of F-hiPSCs present in HipSci: each pair having resulted from the same reprogramming experiment. In all, 54 pairs (46%) of hiPSCs shared more than ten mutations and had similar numbers and profiles of substitutions (cosine similarity >0.9, Fig. 3A). The remaining 64 pairs (54%) of hiPSCs shared ten or fewer substitutions and were dissimilar in mutation burden and profile (cosine similarity <0.9, Fig. 3A). We found some striking differences, for example the F-hiPSC line HPSI0314i-bubh_1, derived from donor HPSI0314i-bubh, had 900 substitutions with no UV signature, whilst HPSI0314i-bubh_3 had 11,000 substitutions with significant representation from UV-associated damage (>90%). Analysis of the parental fibroblast line HPSI0314i-bubh showed some, albeit reduced, evidence of UV-associated mutagenesis (Fig. 3A). Hence, we postulate that F-hiPSCs from the same reprogramming experiment could show considerable variation in mutation burden because of the different levels of sunlight exposure to each skin fibroblast.

**Figure 3.**
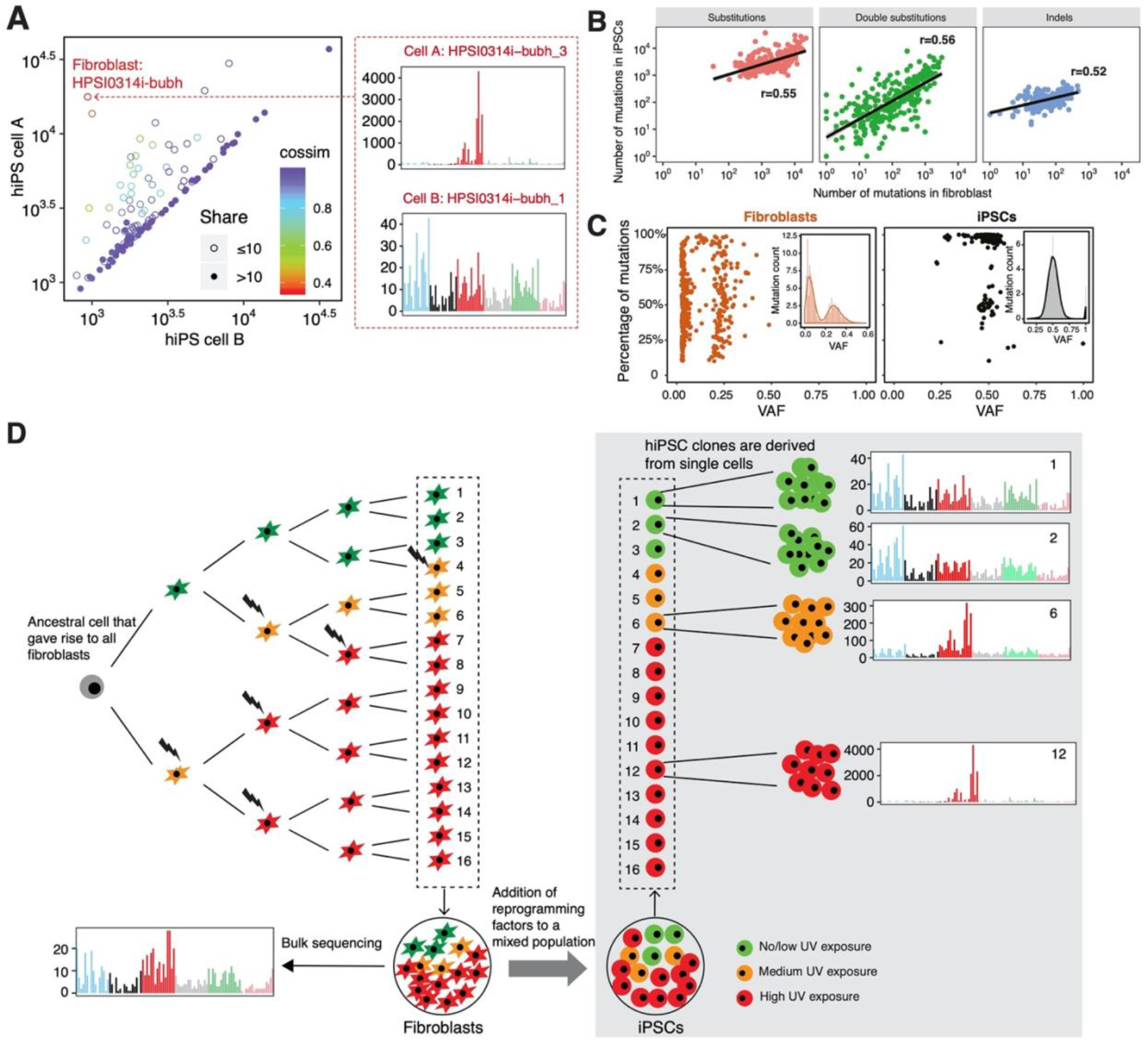
Genomic heterogeneity in F-hiPSCs. (A) Comparison of substitutions carried by pairs of hiPSCs that were derived from fibroblasts in the same reprogramming experiment. Each circle represents a single donor. Hollow circles denote hiPSCs that share fewer than 10 mutations, whilst filled circles indicate hiPSCs that share 10 or more mutations. Colors denote cosine similarity values between mutation profiles of two hiPSCs. Note that cosine similarity is not linear, but a very high score (purple hues) indicates a strong likeness. (B) Correlations between number of mutations in iPSCs and their matched fibroblasts. (C) Summary of subclonal clusters in fibroblasts and hiPSCs. Kernel density estimation was used to smooth the distribution. Local maximums and minimums were calculated to identify subclonal clusters. Each dot represents a cluster which has at least 10% of total mutations in the sample. Most fibroblasts are polyclonal with a cluster VAF near 0.25, whilst hiPSCs are mostly clonal with VAF near 0.5. (D) Schematic illustration of genomic heterogeneity in F-iPSCs. Through bulk sequencing of fibroblasts, all of these cells will carry all the mutations that were present in the grey cell, their most recent common ancestor. The individual cells will also carry their unique mutations depending on the DNA damage received by each cell. Each hiPSC clone is derived from a single cell. Subclone 1 and Subclone 2 cells are more closely-related and could share a lot of mutations in common, because they share a more recent common ancestor. However, they are distinct cells and will create separate hiPSC clones. Subclone 6 (orange cells) and subclone 12 (red cells) are not closely-related to the green cells and have received more DNA damage from UV, making them genomically divergent from the green cells. They could still share some mutations in common because they shared a common ancestor at some early point but will have many of their own unique mutations (largely due to UV damage).

To investigate this further, we analyzed bulk sequenced fibroblasts which revealed high burdens of substitutions, CC>TT double substitutions and indels (Fig. S4A) consistent with UV exposure in the majority of fibroblasts (166/204) (Fig. S4B). Mutation burdens of F-hiPSCs were positively correlated with that of their matched fibroblasts (Fig. 3B) directly implicating the fibroblast population as the root cause of F-hiPSC mutation burden and heterogeneity. Investigating variant allele frequency (VAF) distributions of somatic mutations in F-hiPSCs and fibroblasts demonstrated that most F-hiPSCs were clonal (VAF=0.5), whilst most fibroblasts showed oligo-clonality (VAF<0.5) (Fig. 3C and S5). At least two peaks were observed in VAF distributions of fibroblasts: a peak close to VAF=0 (representing the neutral tail due to accumulation of mutations which follows the power law distribution) and a peak close to VAF=0.25 (representing the subclones in the fibroblast population). From the mean VAF of the cluster, we can estimate the relative size of the subclones, e.g., VAF=0.25 indicating that the subclone occupies half of the fibroblast population. According to this principle, we found that ~68% of fibroblasts contain oligoclonal populations with a VAF<0.25^39^. The mutational burden of F-hiPSCs is thus dependent on which specific cells they were derived from in that oligoclonal fibroblast population. F-hiPSCs derived from the same oligoclonal population will be more similar to each other, whereas hiPSCs derived from different oligoclonal populations within the fibroblast population, could have wildly dissimilar genetic make-ups (Fig. 3D).

There are important implications that arise from the level of subclonal heterogeneity observed in fibroblasts. First, in detection of true mutational damage, it is essential to compare the F-hiPSC genome to a matched germline sample. If instead it is compared to the fibroblast population and the F-hiPSC is derived from a dominant oligoclonal population, many mutations may be detectable in both the fibroblast and F-hiPSC populations and so will be dismissed by mutation-calling software (Fig. S6), giving a false sense of low DNA damage in the F-hiPSC (Fig.S6 and S7). Indeed, we find that ~95% of HipSci F-hiPSCs have shared mutations with matched fibroblasts (Fig. S8). Second, some F-hiPSC mutations may be present in the fibroblasts but not detected through lack of sequencing depth. Comparing WES with high coverage WES (hcWES) data of originating fibroblasts, we found that an increased sequencing depth revealed more coding mutations acquired *in vivo*: WES data showed 47% coding mutations found in hiPSCs were shared with matched fibroblasts, whilst 64% were seen using hcWES data (Fig. S9). The additional hiPSC mutations identified only in hcWES (17%) exhibit a strong UV substitution signature (Fig. S9) suggesting they had been carried over from the fibroblasts. Given recent sequencing studies that have demonstrated a high level of cancer-associated mutations in normal cells^40,41^, it is therefore possible that some driver mutations identified in hiPSCs were acquired *in vivo* and not during cell culture. Third, this work highlights the need for careful clone selection and comprehensive genomic characterization since reprogramming can produce clones with vastly different genetic landscapes.

### Cancer gene mutations in F-hiPSCs

Given the extensive mutagenesis observed in some F-hiPSCs, occasionally reaching thousands per line, we examined all mutations in the coding sequences across the HipSci cohort to determine predicted functional consequences and potential associated selective advantage to F-hiPSCs. We mapped our results to the COSMIC cancer gene census. We found a total of 272 mutations in 145 known cancer genes, across 177 lines (177/452, 39%) from 137 donors. Of note, 151 of 272 mutations were also observed in starting fibroblast populations, even if at lower variant allele fractions, indicating that more than half of these possible “driver” events were acquired *in vivo*. 86 of 272 mutations occurred only once in a Cancer Census gene. It is not clear whether these are definitely driver events although all were scored as “PATHOGENIC” by FATHMM. Nevertheless, 186 of 272 mutations hit 59 genes recurrently in 133 unique lines (133/452, 29%). 38 genes were hit twice (76 variants), while 110 variants occurred in 21 genes with a recurrence of three or more times, involving 77 (17% of lines). All variants that demonstrated gene recurrence three or more times were inspected manually. In particular, we noted recurrent mutations in *BCOR* (11, 2.4%), *CSMD3* (9), *FAT3* (8), *TP53*, *ROS1* and *LRP1B* (7 each), *EP300* and *GRM3* (6 each), *APC* and *PRPTD* (5 each), *CTNND2*, *DCC* and *MED12* (4 each), and *FAT4*, *DNMT3A*, *FGFR1*, *FGFR4*, *KAT6A*, *MAP2K2* and *ZFHX3* (3 each) (Figure 4A, Table S5). Of these, 50/110 (45.5%) were identified in the fibroblast population. In summary, nearly half of cancer-related variants in hiPSC lines that were seen recurrently, had been acquired *in vivo*.

**Figure 4.**
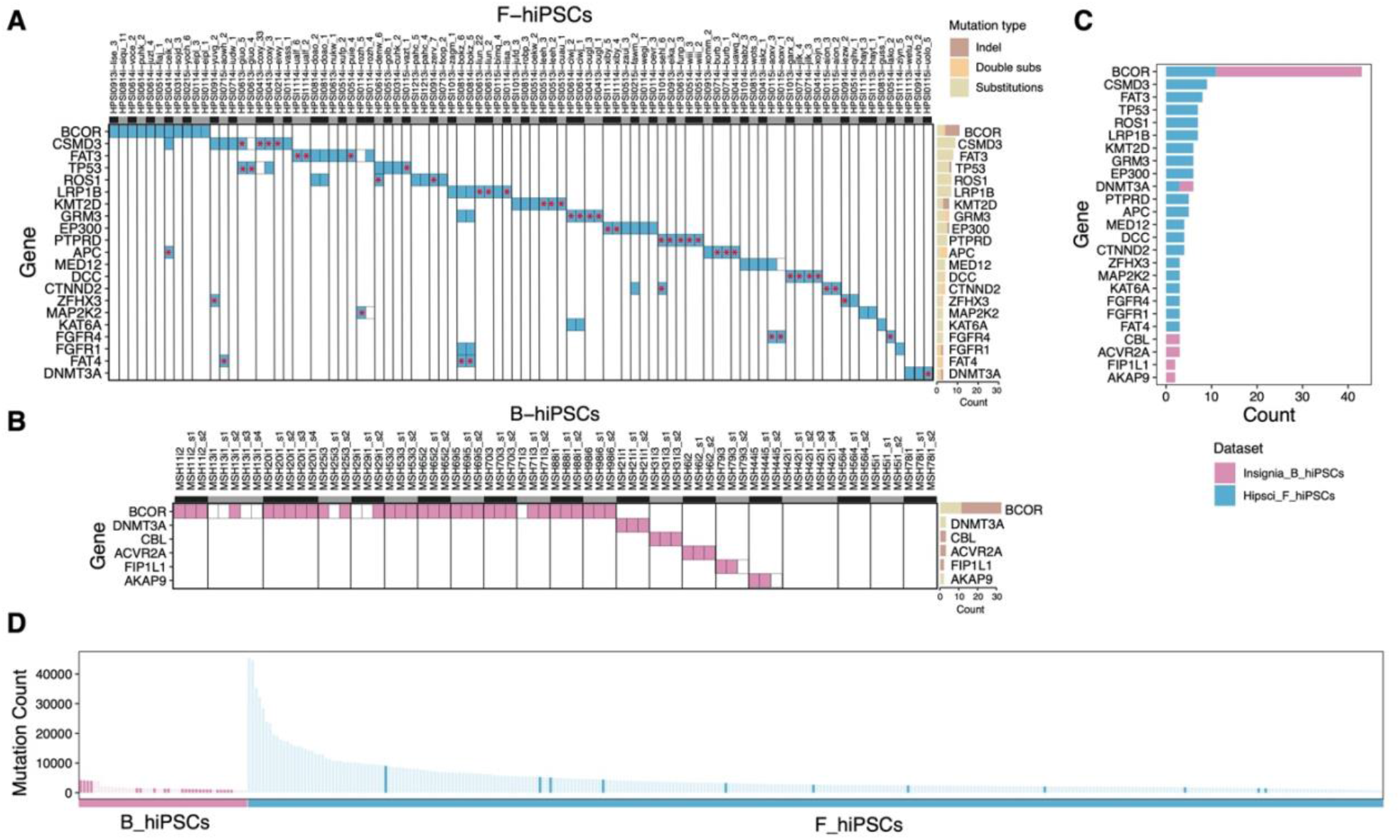
Landscape of recurrently mutated known cancer genes in F-hiPSCs and B-hiPSCs. (A) Recurrent hits in cancer genes observed in F-hiPSC (blue). Red asterisks indicate that the mutations observed in the hiPSC were also observed in their matched corresponding fibroblast samples (i.e. were acquired *in vivo*). Alternating black and grey bars demarcate lines from different donors. 19 donors have two hiPSC lines and 52 donors have single hiPSC lines. Only the donors with lines that had mutations in recurrently hit cancer genes (mutated 3 or more times) are depicted (71). Several hundred lines were inspected but are not depicted in the interest of readability. (B) Hits in cancer genes observed across B-hiPSCs (purple). Right histograms in (A) and (B) summarize the total number of hits per gene. Color indicates the mutation type (substitutions, double-substitutions, indels). (C) Cancer genes with recurrent mutations (>= 3 hits) across B-hiPSCs and F-hiPSCs (D) hiPSCs with BCOR hits are highlighted with a darker shade of purple (B-hiPSCs) or blue (F-hiPSC).

### Strong selection for BCOR mutations in B-hiPSCs

We extended our analysis to B-hiPSCs, derived from erythroblasts, in the HipSci cohort. Intriguingly, three of seventeen sequenced B-hiPSCs carried mutations in *BCOR*, a proportional enrichment when compared to F-hiPSCs in HipSci (p=0.0076, binomial test). However, many B-hiPSCSs in the HipSci cohort did not have matched germline samples to perform subtraction of germline variation, rendering it possible (even if unlikely) for the *BCOR* mutations to be germline in origin. We thus sought alternative cohorts. Blood-derivation methods for generating hiPSCs have been gaining popularity due to the ease of sample collection, but are much less common than F-hiPSCs, and large cohorts of genomically-characterized B-hiPSCs do not exist. Nevertheless, we sought erythroblast-derived hiPSCs created from 21 individuals from the Insignia project on DNA repair abnormalities^42^; 12 patients with inherited DNA repair defects including Oculomotor apraxia type 2 (*AOA2*), ataxia telangiectasia (*ATM*), selenoprotein deficiency (*SECISBP2)*, Lynch Syndrome, Xeroderma Pigmentosum (*XPC*), four patients with exposure to environmental agents (chemotherapy at young age or fetal exposure to maternally-ingested valproate) and five healthy controls. WGS was performed on parental B-hiPSCs and germline DNA of these 21 cases (Table S7). We found 22 mutations in cancer genes including 12 *BCOR* mutations in B-hiPSCs from 12 donors, and one mutation each of *DNMT3A*, *CBL*, *AKAP9, FIP1L1* and *ACVR2A* (Figure 4B). Again, while it is difficult to be certain regarding the selective potential of the single occurrences, the high prevalence of *BCOR* mutations is remarkable. *BCOR* mutations were found in both patients as well as healthy donors.

*BCOR* encodes for BCL6-corepressor protein and is a member of the ankyrin repeat domain containing gene family. The corepressor expressed by *BCOR* binds to BCL6, a DNA-binding protein that acts as a transcription repressor for genes involved in regulation of B cells. Somatic mutations in *BCOR* have been reported in haematological malignancies including acute myeloid leukaemia and myelodysplastic syndromes^43–45^. It has also been reported at a low prevalence in other cancers including lung, endometrial, breast and colon cancers^46^. The high prevalence of *BCOR* mutations in B-hiPSCs but not F-hiPSCs led us to ask whether these could have been derived from haematopoietic stem cell (HSCs) clones that were present in the donors. Clonal haematopoiesis (CH) has been reported in older individuals^47,48^. Some of the commonest genes that are mutated in CH include *DNMT3A, TET2, ASXL1, JAK2* and *TP53*^49–51^. *BCOR* is not one of the most commonly mutated CH genes but has been reported in rare cases of aplastic anemia^45^., although *DNMT3A*, reported once in our cohort, could represent a B-hiPSC derived from a CH clone. Another possibility is that BCOR dysregulation is selected for in the reprogramming/culture process particularly in erythroblast-derived hiPSCs.

### Source of recurrent BCOR mutations

To examine these possibilities, we asked whether the 12 putative selected-for variants could be detected earlier in the B-hiPSC derivation process, either at the erythroblast population stage and/or in the germline DNA sample. We had access to sequences from 21 erythroblast populations and all germline DNAs. However, we did not observe any of these variants in any of the sequencing reads in either erythroblast or germline DNA samples. It is possible that CH clones were not detected due to inadequate sequencing depth.

Notably, five of 12 potential drivers were present at lower VAFs in the B-hiPSCs and could represent subclones within the parental B-hiPSC clone. This would suggest that *BCOR* mutations were arising because of on-going selection pressure in culture, post-erythroblast-transformation and post-reprogramming. To investigate further, we cultured the 21 parental B-hiPSCs for 12-15 days. Two subclones were derived from each of 18 parental B-hiPSCs and four subclones were derived from each of three parental B-hiPSCs, giving a total of 48 subclones assessed by WGS (Figure 4B). Interestingly, five donors had developed new possible driver mutations in B-hiPSC subclones that were not detectable in the parental B-hiPSC population: MSH71 had p.P1229fs*5 *BCOR* mutations in both subclones but not the parent, MSH13 had p.P1115fs*45 *BCOR* mutation in one of four subclones, MSH29 had a p.S1122fs*37 *BCOR* mutation in one of two subclones. MSH42 had a p.F187fs*59 in *POU5F1* in one of two subclones and MSH44 had p.R569H in *LRIG3* in one of two subclones (Figure 4B, Table S8). It is not clear whether the *POU5F1* and *LRIG3* variants are definitely driver events (the latter only reported once in stomach and large intestine cancers previously^52^). Nevertheless, that these possible driver events are present in some but not all subclones and are not present at a detectable frequency in the parental population, suggests that they have arisen late in culture of the parental B-hiPSC. Therefore, in this analysis, we find a high prevalence of recurrent *BCOR* mutations in two independent cohorts of erythroblast-derived hiPSCs (18% of HipSci and 57% of Insignia). We do not find *BCOR* mutations in the originating erythroblasts nor in germline DNA samples. Instead, single-cell-derived subclones variably carried new *BCOR* mutations, suggesting that there may be selection for BCOR dysfunction post-reprogramming in B-hiPSCs. We cannot definitively exclude CH as the source of *BCOR* mutations, as extremely high sequencing depths may be required to detect CH clones. However, *BCOR* is not a particularly common CH mutation and the vast majority of the donors in this cohort were young (<45 yrs) and some were healthy controls, all arguments against *BCOR* being due to CH, and more likely due to selection pressure in B-hiPSCs in culture. That BCOR mutations are seen but at a lower frequency in F-hiPSCs and never in fibroblasts (unlike many of the other cancer genes), hints at a culture-related selection pressure (Figure 4C, D). Why BCOR is so strongly enriched in B-hiPSCs remains unclear. The process of erythroblast-derivation of hiPSCs requires forcefully transforming PBMCs towards the myeloid lineage prior to addition of the Yamanaka factors. Perhaps, this hugely non-physiological process could predispose these cells to recurrent BCOR mutations.

### Long-term culture increases 8-oxo-dG DNA damage

Finally, we examined the genomic effects of *in vitro* hiPSC culture. Mutational signature analyses revealed that all hiPSCs, regardless of primary cell of origin, have imprints of substitution Signature 18, previously hypothesized to be due to oxidative damage in culture^22^ (Fig. 1 and Fig. 2C). Recently, knockouts of *OGG1,* a gene encoding a glycosylase in Base Excision Repair (BER) that specifically removes 8-oxoguanine (8-oxo-dG), have been shown to result in mutational signatures characterized by C>A at A<u>C</u>A>A<u>A</u>A, G<u>C</u>A>G<u>A</u>A, G<u>C</u>T>G<u>A</u>T, identical to Signature 18^53^. Of note, Signature 18 has also been reported in other cell culture systems such as in ES cells^15^, near haploid cell lines^54^ and human tissue organoids in which the contribution to the overall mutation burden was reported to increase with *in vitro* culture^55,56^.

We examined mutations shared between hiPSCs and their matched fibroblasts, representing *in vivo* and/or early *in vitro* mutations, and private mutations that are only present in hiPSCs, representing purely *in vitro* mutations. We observed that Signature 18 is particularly enriched amongst private mutations of nearly all hiPSCs (313 or 97%). By contrast, the majority of hiPSCs (262 or 80%) showed no evidence of this signature amongst shared variants (Fig. 5A, B, C and Fig. S11). We next investigated the relationship between Signature 18 and passage number in the HipSci cohort. We find a positive correlation between Signature 18 and passage number (cor = 0.327; p-value = 5.013e-09, Fig. 5D) reinforcing the notion that prolonged time in culture is likely to be associated with increased acquisition of somatic mutations through elevated levels of DNA damage from 8-oxo-dG (Fig. 5E).

**Figure 5.**
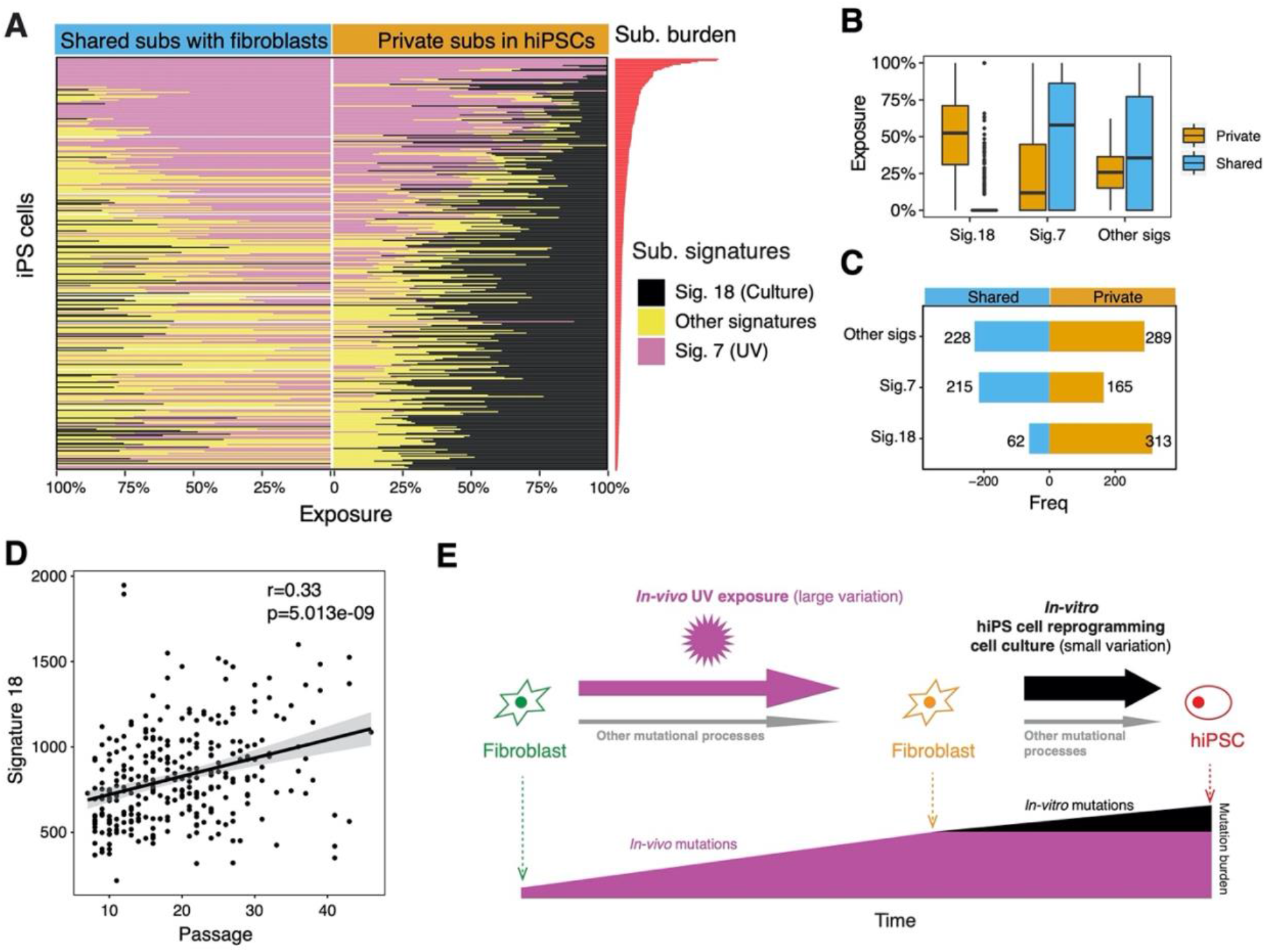
Culture-associated mutagenesis in hiPSCs. (A) Mutations in hiPSCs were separated into two groups: mutations shared between hiPSCs and fibroblasts (left) and mutations that are private to hiPSCs (right). Proportional graphs show substitution signatures of shared and private mutations. Culture-related signature is enriched in private mutations (B) Box-plot of exposures of substitution signatures of shared (blue) and private (orange) mutations. Culture signatures account for ~50% of private substitutions, in contrast to nearly zero amongst shared mutations. Box plots denote median (horizontal line) and 25th to 75th percentiles (boxes). The lower and upper whiskers extend to 1.5× the inter-quartile range (C) Number of samples carrying each signature within shared or private mutations. Most samples acquired the culture signature late in the hiPSC cloning process. (D) Relationship between exposure of culture-related signatures and passage number. (E) Schematic illustration of the mutational processes in somatic fibroblasts and F-hiPSCs.

## Discussion

Human iPS cells are on the verge of entering clinical practice. Therefore, there is a need to better understand the breadth and source of mutations in hiPSCs so as to minimize the risk of harm. Crucially, our work shows that choosing a starting material for pluripotent stem cell derivation may not be straightforward. In an assessment involving 455 cell lines from HipSci, one of the largest hiPSC repositories in the world, we show that whilst skin-derived F-hiPSCs can contain copious UV-associated genomic damage, 39% of F-hiPSCs also carry at least one mutation in a cancer gene. 17% of lines have variants in cancer genes that are recurrently mutated in hiPSCs, although nearly half of these variants are also present in the starting fibroblast material and represent clonal mutations acquired *in vivo*. Notably, the most frequently-mutated gene in F-hiPSCs was *BCOR* and none were observed in corresponding fibroblasts indicating that *BCOR* mutations are likely to have been acquired and selected for *in vitro*. Intriguingly, B-hiPSCs have extraordinary enrichment of *BCOR* mutations, in spite of lower rates of genome-wide mutagenesis and no UV-associated damage. Propagation experiments revealed new *BCOR* mutations arising in some daughter subclones providing support that *BCOR* variants are strongly selected for in hiPSC culture. In addition, both F-hiPSCs and B-hiPSCs exhibit oxidative damage that is increased by long-term culture. Furthermore, following a single reprogramming experiment there can be substantial variation in genomic integrity between different clones. Critically, all of these lines had been previously pre-screened for chromosomal aberrations. Array-based screening techniques whilst useful, do not have the required resolution to report the extent of genomic variation that may be present at nucleotide level. Indeed, further work is required by the community to fully capture the extent and gravity of mutagenesis that could be present in cellular models that are derived using other starting materials too. This work thus highlights potential best practice points to consider when establishing cellular models for clinical or research purposes: We should strive to use a starting cell-type that has minimal genomic damage and to reduce culture time to a minimum. However, comprehensive genomic characterization is indispensable to fully understand the magnitude and significance of mutagenesis in all cellular models.

## Methods

### Samples

The procedures to derive fibroblasts and endothelial progenitor cells (EPCs) from two donors S2 and S7 were previously described^22,57^. Two F-hiPSCs were obtained from S2. Four and two EPC-hiPSCs were obtained from S7 and S2, respectively. 452 F-hiPSCs were obtained from 288 donors and 17 B-hiPSCs were obtained from 9 donors from the HipSci project. 21 B-hiPSCs were obtained from 21 donors from the Insignia project. The details of HipSci and Insignia iPS lines are described below.

### HipSci IPS line generation and growth

All lines were incubated at 37°C and 5% CO234,57. Primary fibroblasts were derived from 2mm punch biopsies collected from organ donors or healthy research volunteers recruited from the NIHR Cambridge BioResource, under ethics for IPS cell derivation (REC 09/H306/73, REC 09/H0304/77-V2 04/01/2013, REC 09/H0304/77-V3 15/03/2013). Biopsy fragments were mechanically dissociated and cultured with fibroblast growth medium (Knockout DMEM with 20% Fetal Bovine Serum (FBS); 10829018, Thermo Fisher Scientific) until outgrowths appeared (within 14 days, on average). Approximately 30 days following dissection, when fibroblasts cultures had reached confluence, the cells were washed with Phosphate Buffered Saline (PBS), passaged using trypsin into a 25 cm^2^ tissue-culture flask, and then again to a 75 cm^2^ flask upon reaching confluence. These cultures were then split into vials for cryopreservation and those seeded for reprogramming, with one frozen vial later used for DNA extraction for whole exome or genome sequencing.

The EPCs were isolated using Ficoll separation of 100mL peripheral blood of organ donors (REC 09/H306/73) and the buffy coat transferred onto a 5μg/cm^2^ collagen (BD Biosciences, 402326) coated T-75 flask. The EPCs were grown using EPC media (EGM-2MV supplemented with growth factors, ascorbic acid plus 20% Hyclone serum; CC-3202, Lonza and HYC-001-331G; Thermo Scientific Hyclone respectively)^57^. EPC colonies appeared after 10 days and these were passaged using trypsin in a 1 in 3 ratio and eventually frozen down using 90% EPC media and 10% DMSO.

Fibroblasts were transduced using Sendai viral vectors expressing human OCT3/4, SOX2, KLF4 and MYC51 (CytoTune, Life Technologies, A1377801) and cultured on irradiated mouse embryonic fibroblasts (MEFs). The EPCs were transduced using four Moloney murine leukemia retroviruses containing the coding sequences of human OCT-4, SOX-2, KLF-4 and C-MYC and also cultured on irradiated MEFs. Following all reprogramming experiments, cells with an iPS cell morphology first appeared approximately 14 to 28 days after transduction, and undifferentiated colonies (six per donor) were picked between day 28 and 40, transferred onto 12-well MEF-CF1 feeder plates and cultured in iPS cell medium with daily medium change until ready to passage. Cells were passaged every five to seven days, depending on the confluence and morphology of the cells, at a maximum 1:3 split ratio until established—usually at passage five or six. Once the iPS cell lines were established in culture, three of the six lines were selected based on morphological qualities (undifferentiated, roundness and compactness of colonies) and expanded for banking and characterization. DNA from fibroblast and iPS cell lines was extracted using Qiagen Chemistry on a QIAcube automated extraction platform.

### Insignia IPS line generation, growth, QC and sequencing

Peripheral blood mononuclear cells (PBMCs) isolation, erythroblast expansion, and IPSC derivation were done by the Cellular Generation and Phenotyping facility at the Wellcome Sanger Institute, Hinxton. Briefly, whole blood samples collected from consented patients were diluted with PBS, and PBMCs were separated using standard Ficoll Paque density gradient centrifugation method. Following the PBMC separation, samples were cultured in media favoring expansion into erythroblasts for 9 days. Reprogramming of erythroblasts enriched fractions was done using non-integrating CytoTune-iPS Sendai Reprogramming kit (Invitrogen) based on the manufacturer’s recommendations. The kit contains three Sendai virus-based reprogramming vectors encoding the four Yamanaka factors, Oct3/4, Sox2, Klf4, and c-Myc. Successful reprogramming was confirmed via genotyping array and expression array. Pluripotency QC is performed based on the QC steps the same as in HipSci project, including the PluriTest using expression microarray data from the Illumina HT12v4 platform and LOH/CNV detection using the HumanExome BeadChip Kit platform. After this, at cell passage equivalent to about 30 doubling, expanded clone were single-cell subcloned to generate 2-4 daughter subclones for each iPSC. Whole-genome sequencing were run on germline, erythroblasts, iPSC parental clone and iPSC subclones.

### Whole exome and genome library preparation and sequencing for HipSci F-hiPSCs

A 96-well plate containing 500ng genomic DNA in 120μl was cherrypicked and an Agilent Bravo robot used to transfer the gDNA into a Covaris plate with glass wells and AFA fibres. This plate was then loaded into the LE220 for the shearing process. The sheared DNA was then transferred out of this plate and into an Eppendorf TwinTec 96 plate using the Agilent Bravo robot. Samples were then purified ready for library prep. In this step, the Agilent NGS Workstation transferred Ampure-XP beads and the sheared DNA to a Nunc deep-well plate, then collected, and washed the bead-bound DNA. The DNA was eluted, and transferred along with the Ampure-XP beads to a fresh Eppendorf TwinTec plate. Library construction comprised of end repair, A-tailing and adapter ligation reactions, performed by a liquid handling robot.

In this step, the Agilent NGS Workstation transferred PEG/NaCl solution and the adapter ligated libraries containing AMPure XP beads to a Nunc deep-well plate, and size selected the bead-bound DNA. The DNA was eluted, and transferred to a fresh Eppendorf TwinTec plate. Agilent Bravo & Labtech Mosquito robotics were used to set up a 384 qPCR plate which was then ready to be assayed on the Roche Lightcycler. The Bravo was used to create a qPCR assay plate. This 384qPCR plate was then placed in the Roche Lightcycler. A Beckman NX08-02 was used to create an equimolar pool of the indexed adapter ligated libraries. The final pool was then assayed against a known set of standards on the ABI StepOne Plus. The data from the qPCR assay was used to determine the concentration of the equimolar pool. The pool was normalised using Beckman NX08-02. All sequencing (paired end) was performed using the Illumina Hiseq platform. The sequencing coverage of WGS, WES and high-coverage WES in hiPSC lines are 41X, 72X and 271X, respectively.

### Sequence alignment, QC, and variant calling for HipSci hiPSCs

Reads were aligned to the human genome assembly GRCh37d5 using bwa version 0.5.1058 (“bwa aln -q 15” and “bwa sampe”) followed by quality score recalibration and indel realignment using GATK version 1.5-959 and duplicate marking using biobambam2 version 0.0.147. VerifyBamID version 1.1.3 was used to check for possible contamination of the cell lines, all but one passed (Fig. S14).

Variable sites were called jointly in each fibroblast and hiPSC sample using BCFtools/mpileup and BCFtools/call version 1.4.25. The initial call set was then pre-filtered to exclude germline variants that were above 0.1% minor allele frequency in 1000 Genomes phase 3^60^ or ExAC 0.3.1^61^. For efficiency we also excluded low coverage sites that cannot reach statistical significance and for subsequent analyses considered only sites that had a minimum sequencing depth of 20 or more reads in both the fibroblast and hiPSC and at least 3 reads with a non-reference allele in either the fibroblast or hiPSC sample. At each variable site we performed a Fisher’s exact test on a two-by-two contingency table, with rows representing the number of reference and alternate reads and the columns the fibroblast or hiPSC sample. We adopted a bespoke approach for mutation calling implemented in BCFtools/ad-bias rather than using existing tools developed for analysis of sequence data from tumor-normal comparisons, because, by definition, tools developed for the analysis of tumor-normal data assume that mutations of interest are absent from the normal tissue. However, in our experiment many mutations were present, albeit at low frequency, in the source tissue fibroblasts. Reversing this, and setting IPS cell lines to be “cancer” and fibroblasts as “normal” introduces an alternative issue, where mutations that are present in IPS cells but absent from fibroblasts are not considered.

More information on bcftools ad-bias can be found on the online bcftools man page: http://samtools.github.io/bcftools/bcftools-man.html. The ad-bias protocol is distributed as a plugin in the main bcftools package, which can be downloaded from http://www.htslib.org/download/. Bcftools ad-bias implements a Fisher test on a 2×2 contingency table that contains read counts of ref / alt alleles found in either the IPS or fibroblast sample. We ran bcftools ad-bias with default settings as follows:

bcftools +ad-bias exome.bcf -- −t1 −s sample.pairs.txt −f ‘%REF\t%ALT\t%CSQ\t%INFO/ExAC\t%INFO/UK1KG’

where “exome.bcf” was the bcf file created by our variant calling pipeline, described in the methods, and sample.pairs.txt was a file that contained matched pairs of the IPSC and corresponding fibroblast sample, one per line as follows:

HPSI0213i-koun_2 HPSI0213pf-koun
HPSI0213i-nawk_55 HPSI0213pf-nawk
HPSI0313i-airc_2 HPSI0313pf-airc
…

We corrected for the total number of tests (84.8M) using the Benjamini-Hochberg procedure at a false discovery rate of 5%, equivalent to a p-value threshold of 9.9×10^−4^, to call a mutation as a significant change in allele frequency between the fibroblast and IPS cell samples. Furthermore, we annotated sites from regions of low mappability and sites that overlapped copy number alterations previously called from array genotypes ^34^ and removed sites that had greater than 0.6 alternate allele frequency in either the fibroblast or hiPSC, as these sites are likely to be enriched for false positives. Dinucleotide mutations were called by sorting mutations occurring in the same IPS cell line by genomic position and marking mutations that were immediately adjacent as dinucleotides.

### Mutation calling of hiPSCs derived from S2, S7 and 10 HipSci lines using fibroblasts as germline controls

Single substitutions were called using CaVEMan (Cancer Variants Through Expectation Maximization; http://cancerit.github.io/CaVEMan/) algorithm^62^. To avoid mapping artefacts, we removed variants with a median alignment score (ASMD) < 90 and those with a clipping index (CLPM) > 0. Indels were called using cgpPindel (http://cancerit.github.io/cgpPindel/). We discarded indels that occurred in repeat regions with repeat count (REP) > 10 and VCF quality (Qual) < 250. Double substitutions were identified as two adjacent single substitutions called by CaVEMan. Ten HipSci lines are: HPSI0714i-iudw_4, HPSI0914i-laey_4, HPSI0114i-eipl_1, HPSI0414i-oaqd_2, HPSI0414i-oaqd_3, HPSI1014i-quls_2, HPSI1013i-yemz_3, HPSI0614i-paab_3, HPSI1113i-qorq_2 and HPSI0215i-fawm_4.

### Mutation calling of Insignia B-hiPSCs using blood as germline controls

Single substitutions were called using CaVEMan (Cancer Variants Through Expectation Maximization; http://cancerit.github.io/CaVEMan/) algorithm^62^. To avoid mapping artefacts, we removed variants with a median alignment score (ASMD) < 90 and those with a clipping index (CLPM) > 0. Indels were called using cgpPindel (http://cancerit.github.io/cgpPindel/). We discarded indels that occurred in repeat regions with repeat count (REP) > 10 and VCF quality (Qual) < 250. Double substitutions were identified as two adjacent single substitutions called by CaVEMan. Mutation calls were obtained for erythroblasts, iPSC parental clones and subclones.

### Mutational signature analysis

Mutational signature analysis was performed on the WGS data set. All dinucleotide mutations were excluded from this analysis. We generated 96-channel single substitution profile for 324 hiPSCs and 204 fibroblasts. We fitted previous discovered skin-specific substitution to each sample using an R package (signature.tools.lib)^31^. Function SignatureFit_withBootstrap() was used with default parameters. In downstream analysis, the exposure of two UV-caused signatures Skin_D and Skin_J were summed up to represent the total signature exposure caused by UV (Signature 7). A *de novo* signature extraction was performed on 324 skin-derived WGS hiPSCs to confirm that the UV-associated skin signatures (Skin_D and Skin_J, Signature 7) and culture-associated one (Skin_A, Signature 18) are also the most prominent signatures identified in de novo signature extraction (Fig. S13).

### Analysis of C>T/CC>>TT transcriptional strand bias in replication timing regions

Reference information of replication-timing regions were obtained from Repli-seq data of the ENCODE project (https://www.encodeproject.org/)^63^. The transcriptional strand coordinates were inferred from the known footprints and transcriptional direction of protein coding genes. In our dataset, we first orientated all G>A and GG>AA to C>T and CC>TT (using pyrimidine as the mutated base). Then we mapped C>T and CC>TT to the genomic coordinates of all gene footprints and replication timing regions. Lastly, we counted the number of C>T/CC>TT mutations on transcribed and untranscribed gene regions in different replication timing regions.

### Identification of fibroblast-shared mutations and private mutations in hiPSCs

We classified mutations (substitutions and indels) in hiPSCs into fibroblast-shared mutations and private mutations. Fibroblast-shared mutations in hiPSC are the ones that have at least one read from the mutant allele found in the corresponding fibroblast. Private mutations are the ones that have no reads from the mutant allele in the fibroblast. Mutational signature fitting was performed separately for fibroblast-shared substitutions and private substitutions in hiPSCs. For indels, only the percentage of different indel types were compared between fibroblast-shared indels and private indels.

### Clonality of samples

We inspected the distribution of VAFs of substitutions in fibroblasts and hiPSCs. Almost all hiPSCs had VAFs distributed around 50%, indicating that they were clonal. In contrast, all fibroblasts had lower VAFs, which distributed around 25% or lower, indicating that they were oligoclonal. We computed kernel density estimates for VAF distributions of each sample. Based on the kernel density estimation, the number of clusters in a VAF distribution was determined by identification of the local maximum. Accordingly, the size of each cluster was estimated by summing up mutations having VAF between two local minimums.

### Variant consequence annotation

Variant consequences were calculated using the Variant Effect Predictor^64^ and BCFtools/csq^65^. For dinucleotide mutations, we recorded only the most impactful consequence of other of the two members of the dinucleotide, where the scale from least to most impactful was: intergenic, intronic, synonymous, 3’ UTR, 5’ UTR, splice region, missense, splice donor, splice acceptor, start lost, stop lost and stop gained. We identified overlaps with putative cancer driver mutations using the COSMIC “All Mutations in Census Genes” mutation list (CosmicMutantExportCensus.tsv.gz) version 92, 27th August 2020. We further filtered Mutations with FATHMM scores < 0.7 from driver analysis, as pathogenic mutations require FATHMM scores ≥ 0.7^66^.

### Other statistical analysis

All statistical analysis were performed in R^67^. The effects of age and sex on mutation burden of F-hiPSCs were estimated using Mann-Whitney test, wilcox.test() in R. Tests for correlation in the study were performed using cor.test() in R. A simple linear model was fitted to calculate the coefficients of determination (R-squared) between mutational signatures and different mutation types.

## Supporting information

Table_S2_skin_blood_subs_indels_burden

Table_S3_HipSci_sample_summary_3experiments

Table_S4_ips_summary_muts

Table_S5_hipsci_drivers

Table_S7_Insignia_ips

Table_S8_Insignia_pathogenic_muts

## Data availability

Links to raw sequencing data produced in this study are available from the HipSci project website (www.hipsci.org). Raw data are deposited in the European Nucleotide Archive (ENA) under the following accessions: ERP006946 (WES, open access samples) and ERP017015 (WGS, open access samples), and in the European Genotype-Phenotype Archive under accession EGAS00001000592 (WES, managed access samples). The open access data samples are freely available to download, while the managed access data are available following a request and a data access agreement. Variant call sets, summary files and code will be made available on publication.

## Acknowledgments

We thank the subjects and volunteers who agreed to participate in the studies.

## Author contributions

FR, XZ, PD, DG and SNZ conceived this study, designed the experiments and wrote the manuscript. RD contributed to the methodology of the study. XZ, TDA, GK, QW, YM and IM performed bioinformatic analysis of the sequencing data. ARB reviewed and edited the manuscript.

## Extended data

Supplementary Information. Figures S1-S13.

Table S2. Summary of mutation burden for samples in Figure 1.

Table S3. Summary of samples from HipSci project.

Table S4. Mutation burden of F-hiPSCs from HipSci project.

Table S5. Cancer driver mutations identified in HipSci F-hiPSCs.

Table S6. Somatic and germline mutations used in functional analysis.

Table S7. Mutation burden of B-hiPSCs from Insignia project.

Table S8. Cancer driver mutations identified in Insignia B-hiPSCs.

