## Supplementary figures for "Substantial somatic genomic variation and selection for *BCOR* mutations in human induced pluripotent stem cells"

**Supplementary Table**

| | **Donors** | **Parental lines** | **Subclone lines** | **Sequencing method** | **"normal" used to remove germline mutations** |
|---|---|---|---|---|---|
| **F-hiPSCs** | S2 | 2 | 0 | WGS | Fibroblast |
| | 9 from HipSci | 10 | 0 | WGS | Fibroblast |
| | HipSci 288 healthy donors | 452 | 0 | 324 WGS<br>381 WES<br>106hcWES | Bespoke approach |
| **B-hiPSCs** | S2 | 2 | 0 | WGS | Fibroblast |
| | S7 | 4 | 0 | WGS | Fibroblast |
| | Insignia 21 patients | 21 | 48 | WGS | Germline control |
| | HipSci 9 donors | 17 | 0 | WES | - |

Table S1. Summary of hiPSC samples.

### Supplementary Figures

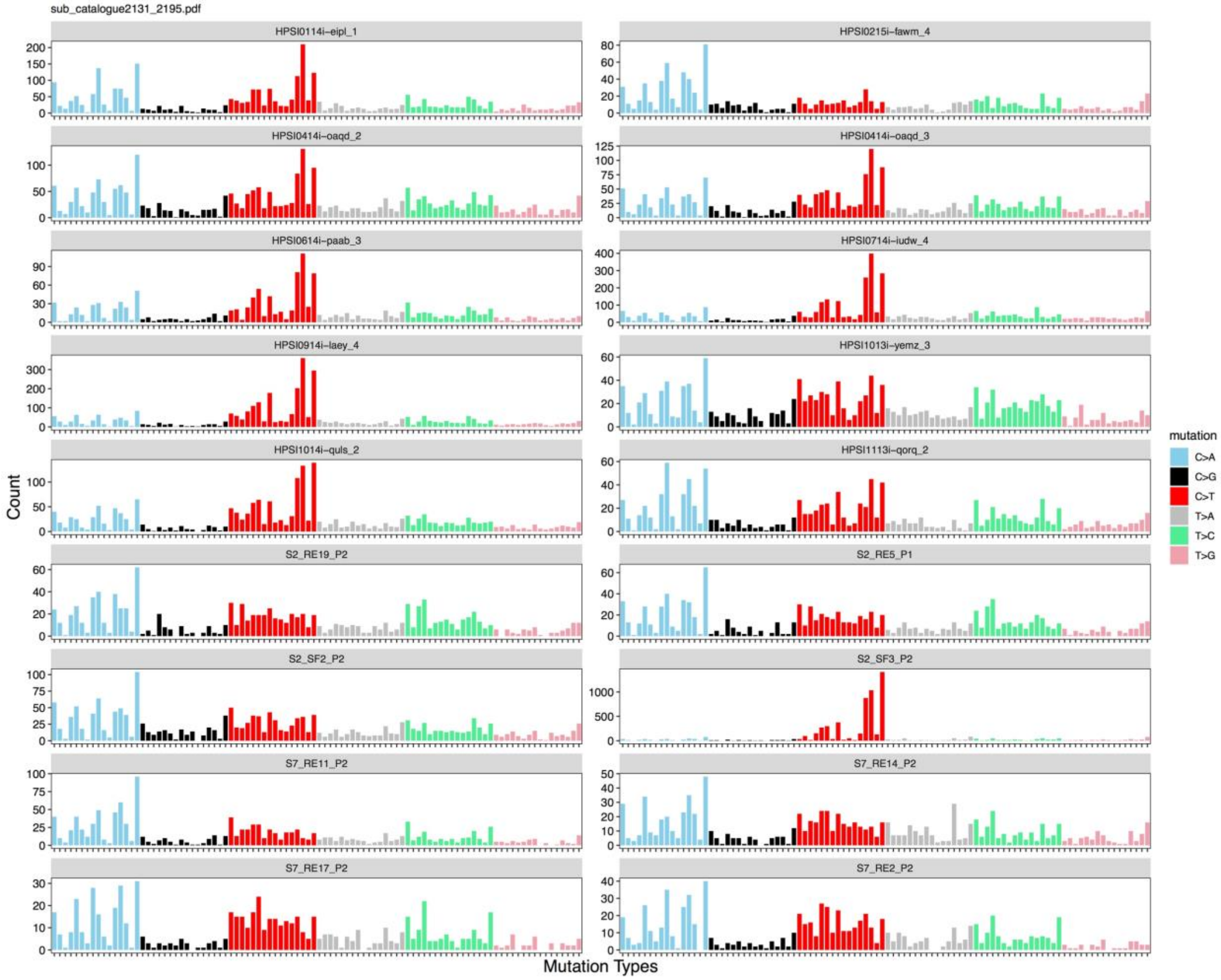


Figure S1. Mutational profiles of 18 blood-derived iPSCs and skin-derived iPSCs featured in Figure 1.

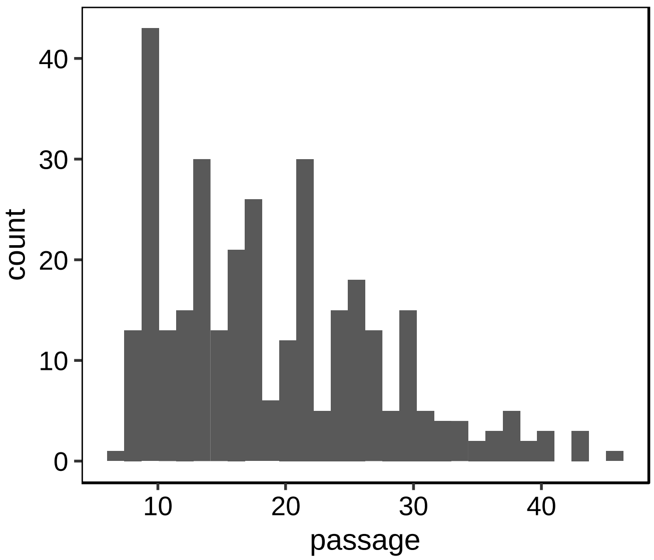


Figure S2. Distribution of passage number of iPSCs in HipSci. The median is 18.

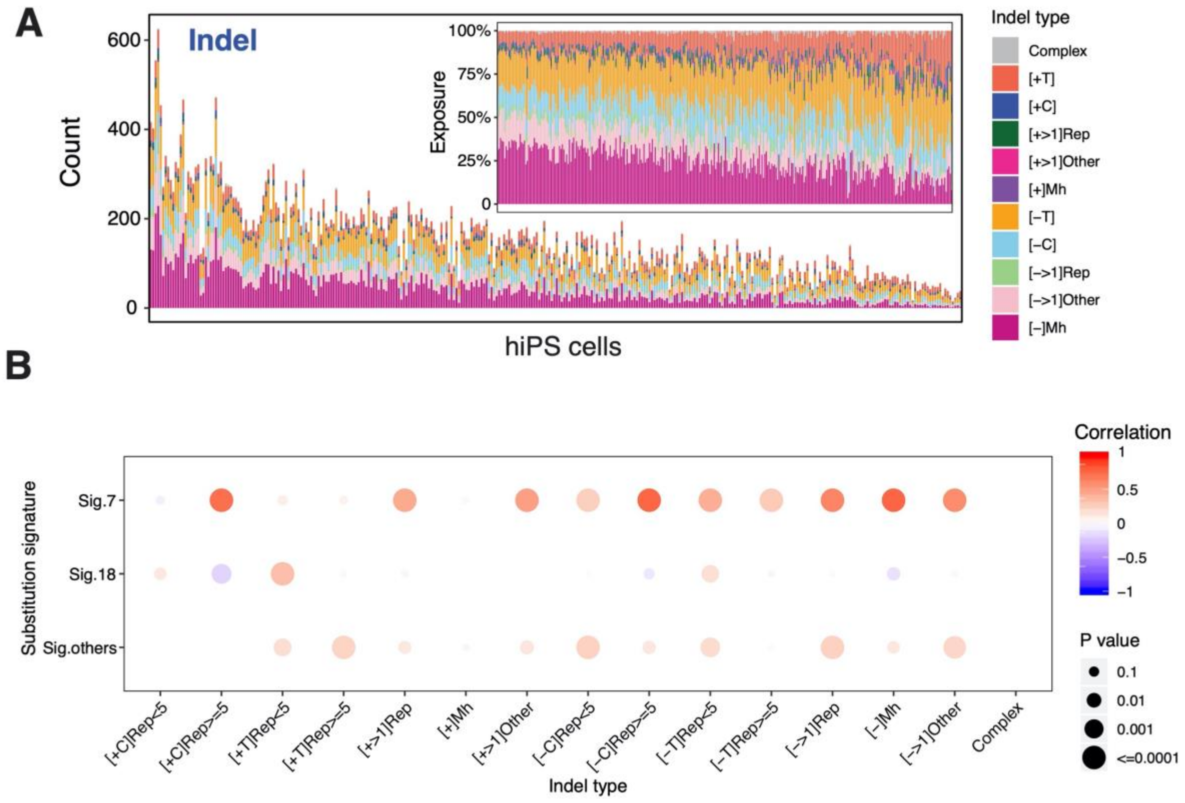


Figure S3. (A) Distribution of mutational signatures in 324 fibroblast-derived iPSC lines. The inset figure shows the relative exposures of mutational signatures/indel types. (B) Correlation between substitution signatures and indel types.

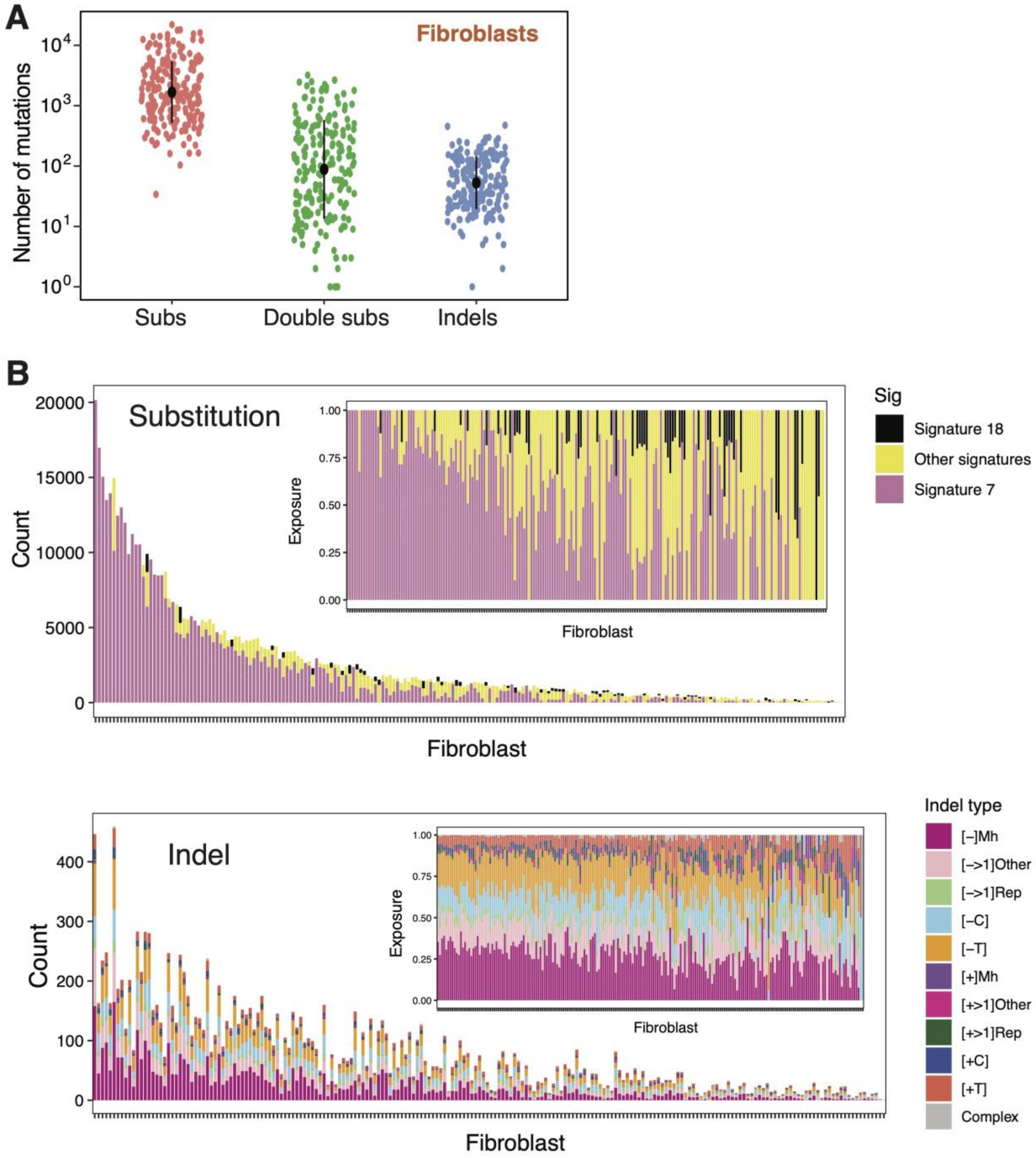


Figure S4. Mutation burden and mutational signatures in fibroblasts. (A) Mutation burden of substitutions, CC>TT double substitutions and indels in fibroblasts. (B) The amount of each mutational signature and indel type (exposure) in fibroblasts.

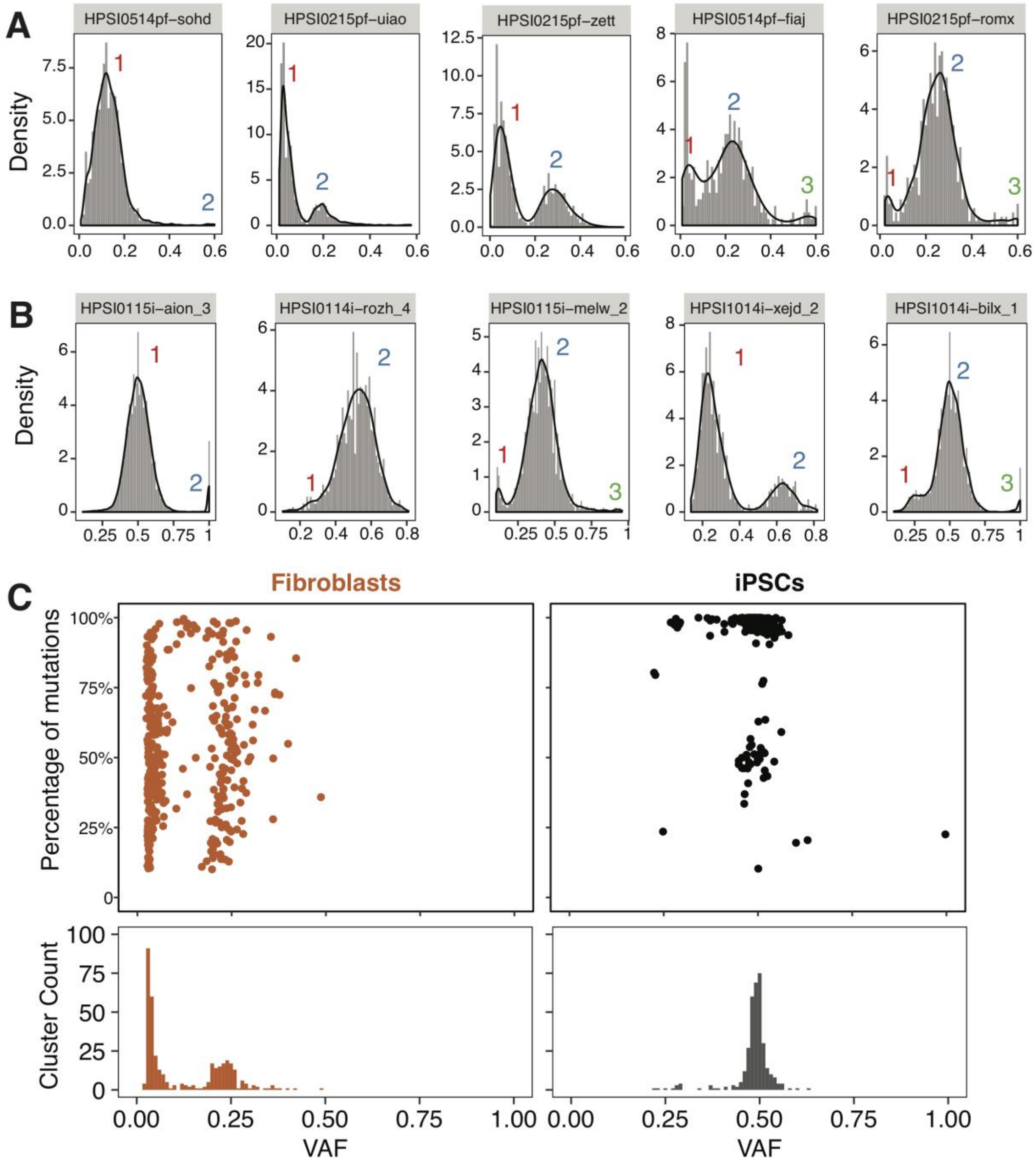


Figure S5. Analysis of variant allele frequency in fibroblasts and iPSCs. Distribution of variant allele frequency distribution of five fibroblasts and five hiPSCs are shown in (A) and (B), respectively. Kernel density estimation was used to smooth the distribution. Local maximums and minimums were calculated to identify subclonal clusters. (C) Summary of subclonal clusters in fibroblasts and hiPSCs with. Each dot represents a cluster which has at least 10% of total mutations in the sample. Most of fibroblasts are polyclonal with VAF of a cluster near 0.25, whilst hiPSCs are mostly clonal with VAF near 0.5.

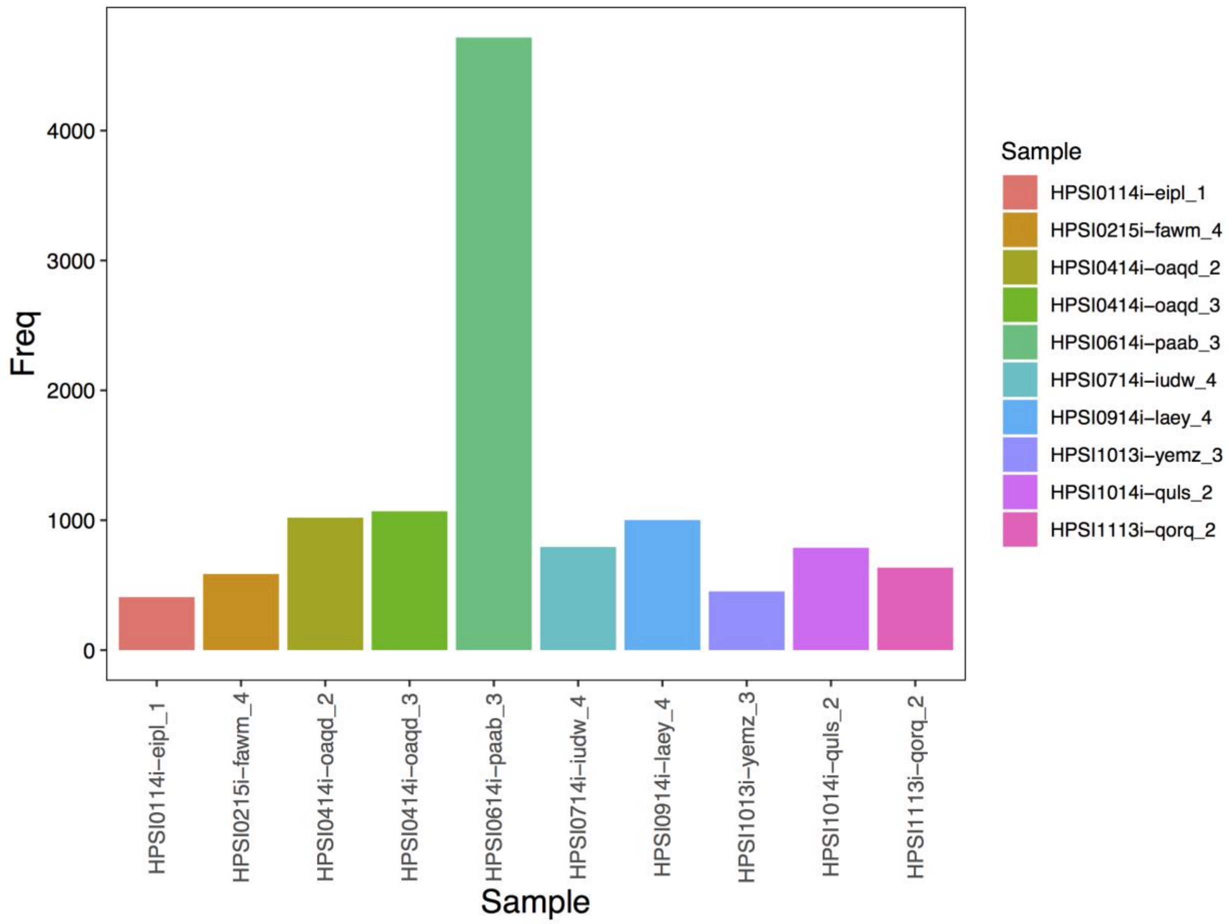


Figure S6. Number of substitutions that were removed from iPSCs using fibroblast as "normal" for ten HipSci samples from Figure 1.

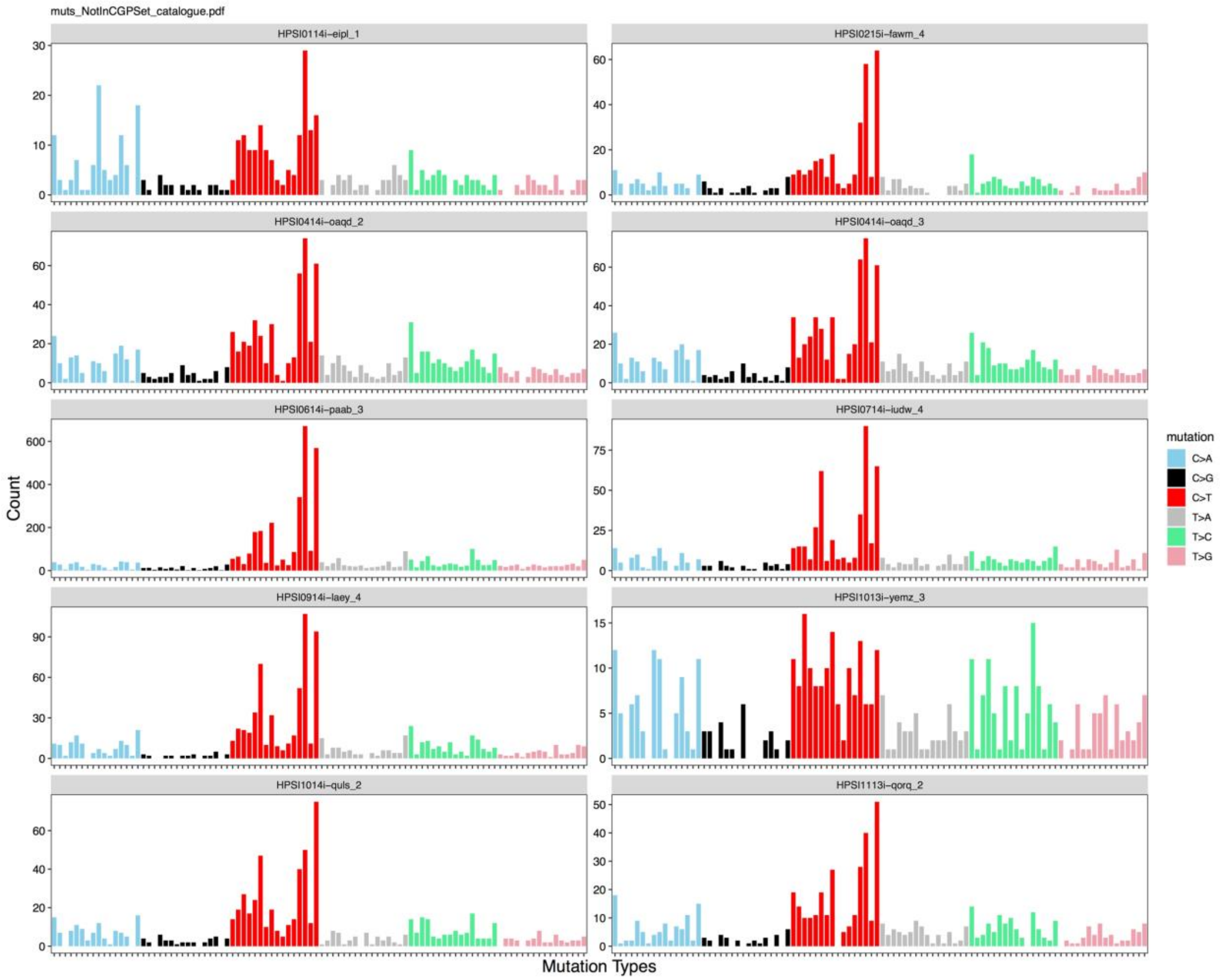


Figure S7. Mutation profile of substitutions that were removed from iPSCs using fibroblast as “normal” for ten HipSci samples from Figure 1. These removed mutations are not germline SNPs, and mostly composed of mutations that are typical of UV exposure.

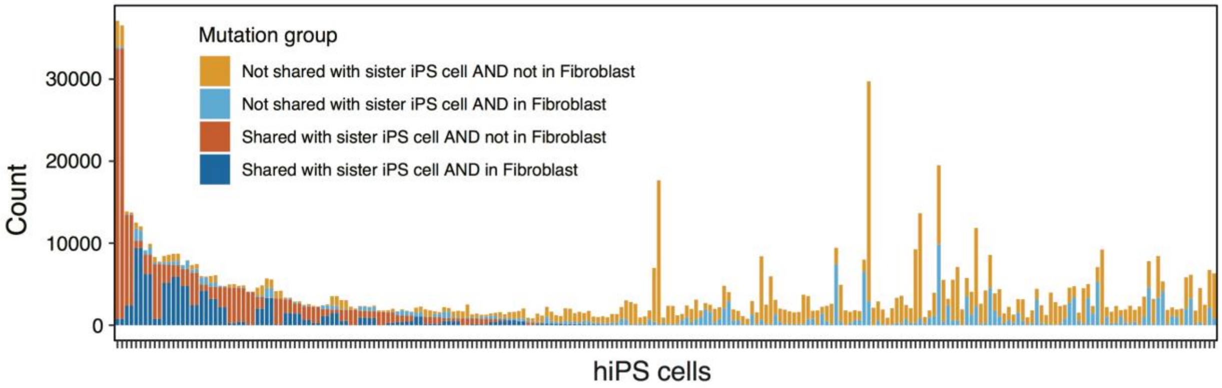

Figure S8. Shared mutations between hiPSCs and the matched fibroblasts.

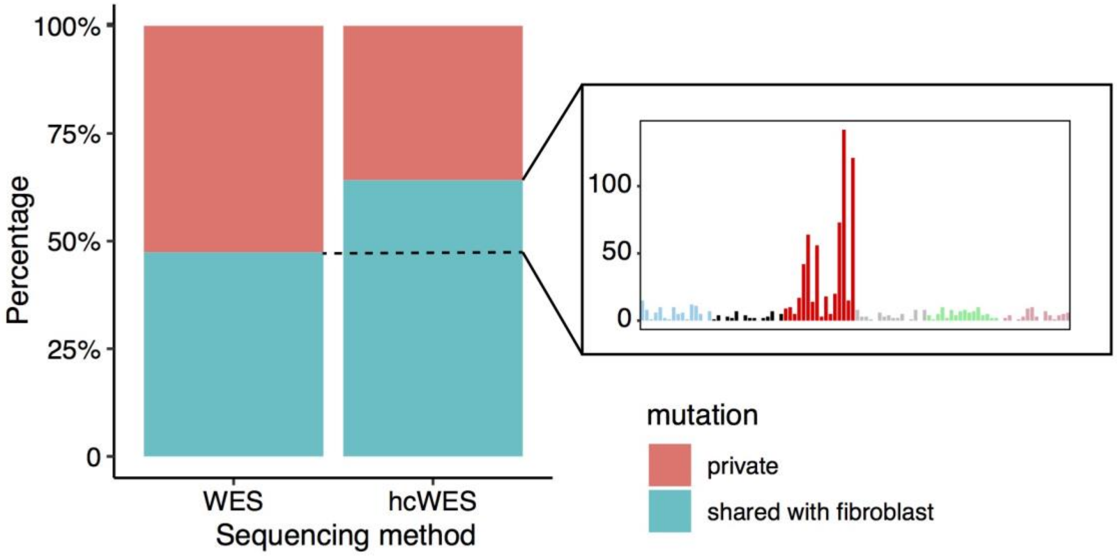

Figure S9. Comparison of WES (72X) and high coverage WES (hcWES, 271X) of fibroblasts. More mutations in hiPSCs were discovered in fibroblasts (shared mutations) through hcWES than through WES, resulting in the percentage of shared-mutations increased in hcWES data. Interestingly, the mutational profile of these increased shared-mutations is very similar to the UV signature, indicating that increasing sequencing depth enables more UV-caused somatic mutations that were found in hiPSCs to also be detected in the corresponding fibroblasts.

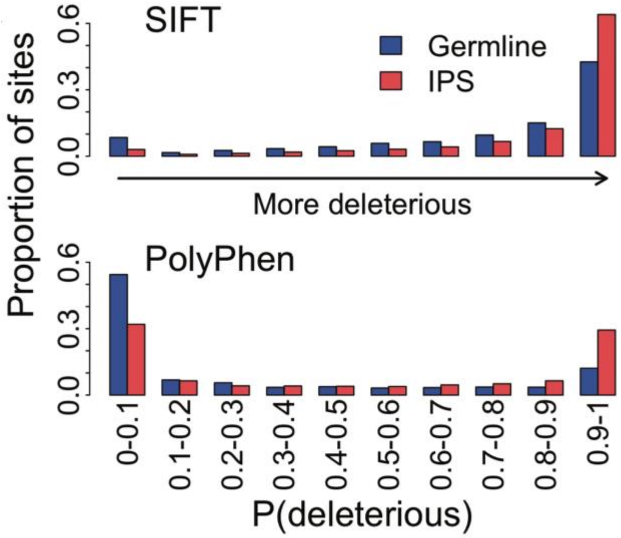


Figure S10. Distribution of the probability of being deleterious (P(deleterious)): either 1-SIFT score (top panel) or the PolyPhen2 score (bottom panel) in IPS mutated sites and in germline (AF > 1% in UK10K and ExAC).

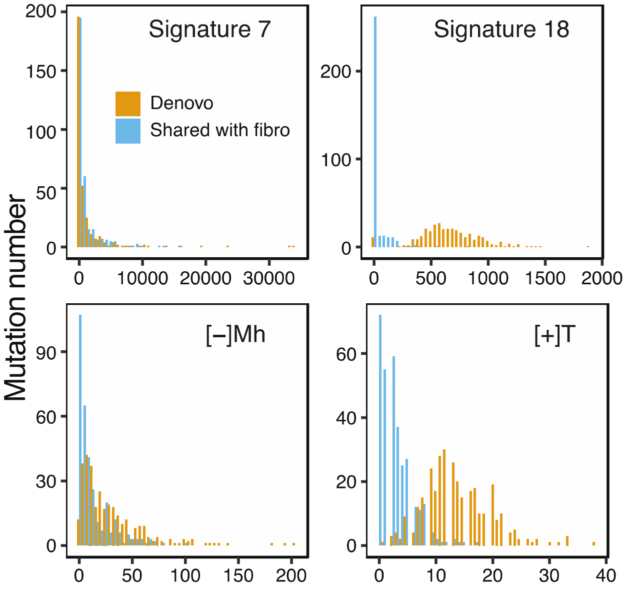


Figure S11. Histogram of shared and private mutations for signature 7 (UV), signature 18 (oxidative damage), [-]Mh and [+]T.

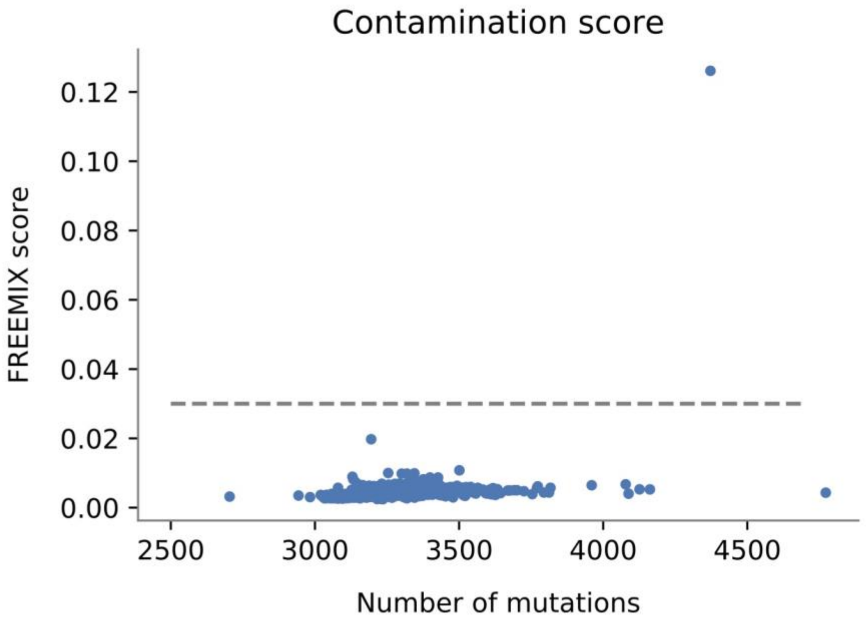


Figure S12. There is no evidence of contamination except for one cell line and there is no correlation between the number of mutations and the FREEMIX score ($R^2$=0.1). The dashed line at 0.03 is the threshold suggested by VerifyBamID to accept or potentially flag the sample as contaminated. The outlier cell line (HPSI0913pf-coyi) was removed from analysis.

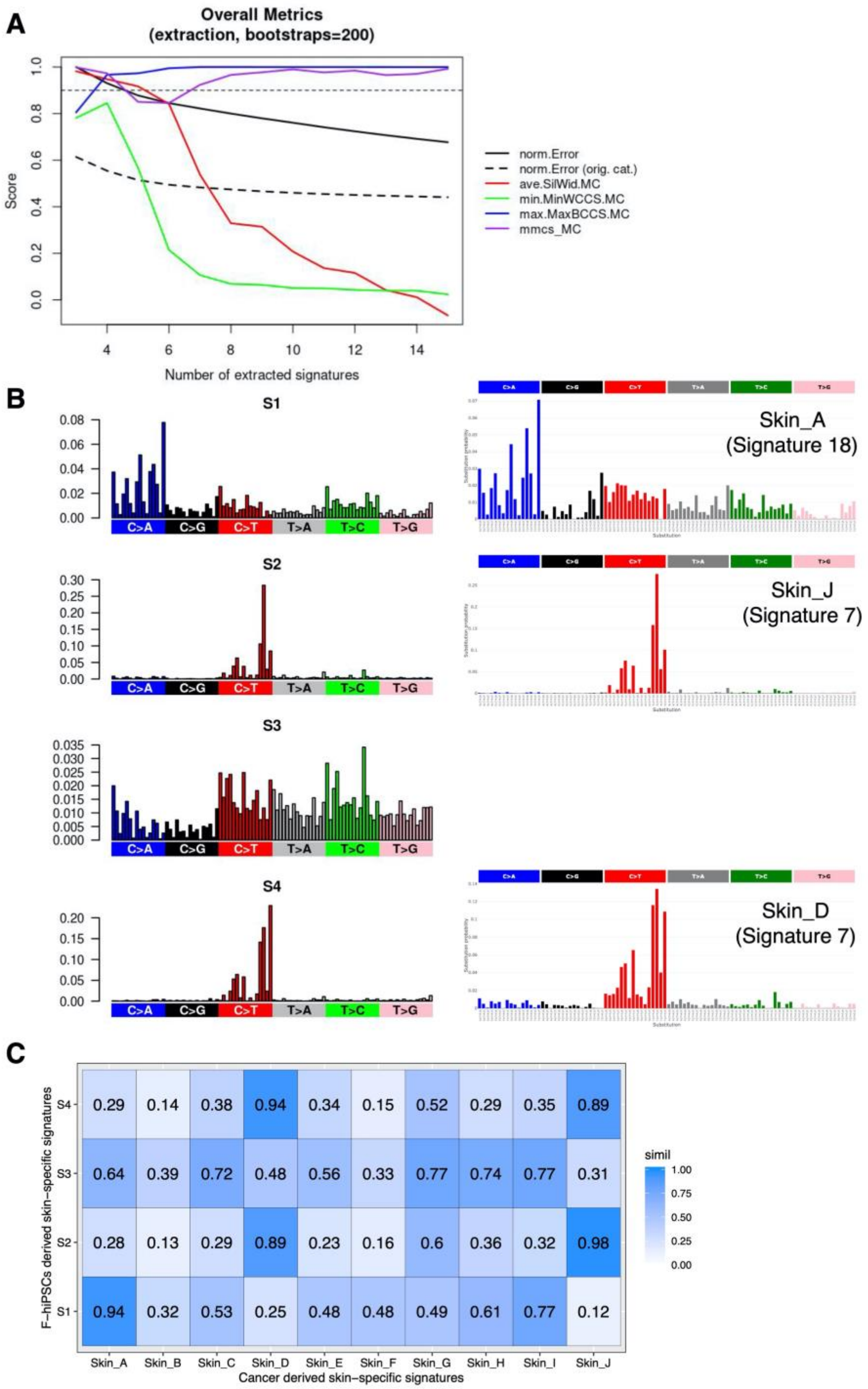


Figure S13. *De novo* extraction on 324 skin-derived WGS hiPSCs from the HipSci project. (A) Metrics for selecting optimal number of signatures. (B) Four mutational signatures extracted from this data set. Profile of similar skin cancer derived signatures are shown. (C) Cosine similarities between F-iPSCs signatures and skin cancer derived signatures. S2 and S4 are most similar (cossim: 0.94-0.98) to UV-associated mutational signatures, Skin_J and Skin_D (signature 7), respectively. S1 is most similar to Skin_A (signature 18), the culture signature (cossim: 0.94). S3 does not show high similarity to any skinspecific signatures (cossim <0.8), but also has very low probabilities for all 96 channels (note y-axis values are very small), and the relatively featureless profile would suggest that it is likely to be “noise”. This is not uncommon in signature extractions.
